# Firefighter Helmets and Cervical Intervertebral Kinematics: An OpenSim-Based Biomechanical Study

**DOI:** 10.1101/2023.11.16.567468

**Authors:** Gustavo M. Paulon, S. Sudeesh, Suman K. Chowdhury

## Abstract

The assessment of cervical intervertebral kinematics can serve as the basis for understanding any degenerative changes in the cervical spine due to the prolonged wear of a heavyweight, imbalanced firefighting helmet. Therefore, this study aimed to analyze cervical intervertebral kinematics using OpenSim musculoskeletal modeling platform in order to provide much-needed insights of how the inertial properties of firefighter helmet affect cervical spinal mobility. A total of 36 firefighters (18 males and 18 females) were recruited to perform static and dynamic neck flexion, extension, and left and right lateral bending tasks for three conditions: 1) no-helmet, 2) US-style helmet with a comparatively superior center of mass (COM), and 3) European-style helmet with relatively higher mass but an inferior COM. Three custom-made OpenSim head-neck models were created to calculate cervical intervertebral kinematics for each helmet condition. Results showed that the helmet use significantly (p<0.001) affects neck and cervical spinal kinematics. Especially, the superior COM placement in the US-style helmet, despite its lighter weight, caused more pronounced kinematic changes and quicker attainment of peak flexion and extension angles compared to the European-style helmet across all cervical joints. Moreover, results also revealed discrepancies between OpenSim-derived neck and cervical range-of-motion and those reported in previous in-vivo studies. In conclusion, the present study underscores the importance of designing firefighter helmets with a lower profile (less superior COM) to enhance neck range of motion and minimize potential neck injuries.

## Introduction

Neck pain is a multifactorial musculoskeletal health condition with a high global prevalence. According to the 2019 Global Burden of Disease (GBD) Study, neck pain was among the top four most prevalent musculoskeletal disorders (MSDs) and affected 222.7 million people (2.9% of total population) worldwide (GBD, 2019). Various factors, ranging from ergonomic factors in the workplace to daily lifestyle and recreational activities, contribute to the onset and persistence of neck pain. For instances, prolonged static postures (Christensen et al., 2023), repetitive neck motions (Guidotti, 1992), heavy lifting or working in awkward neck postures (Ariens et al., 2000), sedentary lifestyle and poor sleep quality (Peterson and Pihlström, 2021), and chronic stress and certain mental health conditions (Kim et al., 2013) have been associated with work-related neck MSDs. In addition to these causes, the usage of helmets and other head mounted devices in medical surgery (Nimbarte et al., 2013), military (Hanks et al., 2018), firefighting (Park et al., 2014; Wang et al., 2021), sports (Kent et al., 2020), and vehicle safety applications (Diyana et al., 2019) were found to increase the likelihood of neck MSDs.

Previous biomechanical studies have primarily associated excessive weight and shifted center of mass (COM) of helmets with the risks of neck injuries. For instance, Van Dijke et al. (1993) and Newman et al. (2022) studied the influence of jet pilot helmets’ inertial properties on the risk of neck injury and found that helmets significantly increase the neck joint reaction moments. Barrett et al. (2023) showed that the inertial properties (i.e., COM, mass, and moment of inertia) of a military helmet resulted in higher compressive forces in the cervical spine. Other studies on construction (Boschman et al., 2015), mining (Torma-Krajewski et al., 2006), and firefighting (Wang et al., 2021) helmets also emphasized the importance of enhancing the ergonomic aspects (i.e., inertial properties) of helmets in order to reduce neck discomfort and MSDs. Among them, firefighter helmets are usually heavier than other helmet types because they include two shells and other padding materials to provide high flame and impact protection to the users. Additionally, the modern firefighter helmets include other supporting accessories, such as face-shield, visor, and lighting equipment. Their additions can further increase the likelihood of cervical spinal injuries due to the increase in the total helmet weight and a potential shift of the helmet COM. Especially, these changes can make female users more vulnerable to neck pain as they have weaker neck muscles (Vasavada et al., 2008) and smaller cervical vertebra (Stemper et al., 2008) than males. It was also reported that about 23 % of all firefighter injuries (27,150 cases out of 118,070 cases) are related to the head, neck, and shoulder – out of which, 5.5% involved strains or overexertion on the head and neck (Campbell and Molis, 2022). As firefighter helmets are worn daily for a prolonged duration, their repetitive usages over time can result in degenerative neck injuries, such as spinal cord spondylosis, degenerative disc diseases, and ossification of the ligamentum flavum (Echarri and Forriol, 2005).

As indicated in previous clinical (Hino et al., 1999; Hirsch et al., 1967) and biomechanical (Panjabi et al., 2001) studies, the assessment of cervical intervertebral kinematics can be used as surrogate measures of degenerative changes occurring in the cervical spine because these changes can adversely affect the neck joint mobility. Thus, an accurate information about the cervical intervertebral joint mobility under the effects of no-helmet and helmet conditions can serve as the basis to evaluate how the helmet inertial properties may cause abnormal neck conditions. A number of in-vivo experimental studies using dynamic fluoroscopic system (Zhou et al., 2020), dynamic stereo-radiographic system (Anderst et al., 2015b), and three-dimensional MRI (Ishii et al., 2006) have reported neck and cervical intervertebral ROM. As these in-vivo studies are labor-, time-, and technology-intensive, a handful of studies used in-silico methods, such as OpenSim (an open-source musculoskeletal modeling platform) (Barrett et al., 2022a; Barrett et al., 2022b; Mathys and Ferguson, 2012; Newman et al., 2022) because of their non-invasiveness nature and ease-of-use in modeling the head-helmet dynamics. Nevertheless, they overlooked the influence of helmet on cervical intervertebral kinematics, especially during dynamic neck movements. Additionally, our systematic literature review showed only one survey-based study on the firefighter helmet by Wang et al. (2021). They reported that a poorly-fitting, imbalanced helmet can restrict firefighter’s neck mobility, which motivates us to conduct a thorough biomechanical investigation to assess the impacts of firefighter helmets on cervical intervertebral kinematics — the potential surrogate measures of degenerative neck injuries.

Therefore, this study aimed to investigate the effects of firefighter helmet inertial properties on neck and cervical intervertebral kinematics, especially during various neck dynamic movements. Two types of helmets are frequently used by firefighters: 1) US-style helmet (resemblance to hats) and 2) European-style helmet (resemblance to aviation helmets). A quantitative analysis of their inertial properties and how their variations affect cervical intervertebral kinematics would provide an unprecedented understanding of firefighter’s neck MSD mechanisms and assist practitioners to design an injury-mitigating firefighter helmet that can be worn for prolonged time.

## Methods

### Participants

We recruited eighteen male (weight: 88.5 ± 18.9 kg; height: 1.77 ± 0.09 m; BMI: 28.8 ± 5.39; Age: 39.2 ± 7.23 years) and eighteen female (weight: 69.5 ± 12.2 kg; height: 1.64 ± 0.05 m; BMI: 24.2 ± 2.85; Age: 31.2 ± 8.62 years) firefighters from the local fire departments. The inclusion criteria required all participants to be healthy and did not have any recent history of neck, shoulder, and back injury. Prior to their participation, they signed an informed consent form approved by the local Institutional Review Board (IRB2020-708).

### Experiment

Participants primarily performed two repetitions of static task—holding a static head-neck position in full flexion, extension, and left and right lateral bending for five seconds—and two repetitions of dynamic task—self-paced, controlled (using a digital metronome) full flexion-extension and left-right lateral bending movements, each separately for 3 seconds— using three different helmet conditions: 1) baseline, no-helmet, 2) US-style helmet (Bullard UM6WH), and 3) European-style (Cairns XF1) helmet in a random order (Figure 1). We collected full-body movements using a 10-camera motion capture system at 60 Hz (Krestel 1300; Motion Analysis, California, USA) and ground reaction forces using two force plates (Bertec, Ohio, USA) at 600 Hz. We also used a handheld, rotating 3D scanner (EinScan HX, Shining 3D, Hangzhou, China) to image individual helmets at 20 Hz frame rate. The kinematics data were pre-processed in Cortex-9 software (Motion Analysis Corporation, Rohnert Park, California, USA). Though we used full-body plug-in gait marker set (Vicon, 2023) consisting of 44 markers in the experimental protocol, we created additional 32 virtual markers in Cortex to measure link-segment anthropometric measures accurately. The force plate data were exported from Cortex-9 as C3D format and then converted to motion (.mot) files using C3Dtools (Mokhtarzadeh and Bagheri, 2023) in order to input them into OpenSim.

**Figure 1:**
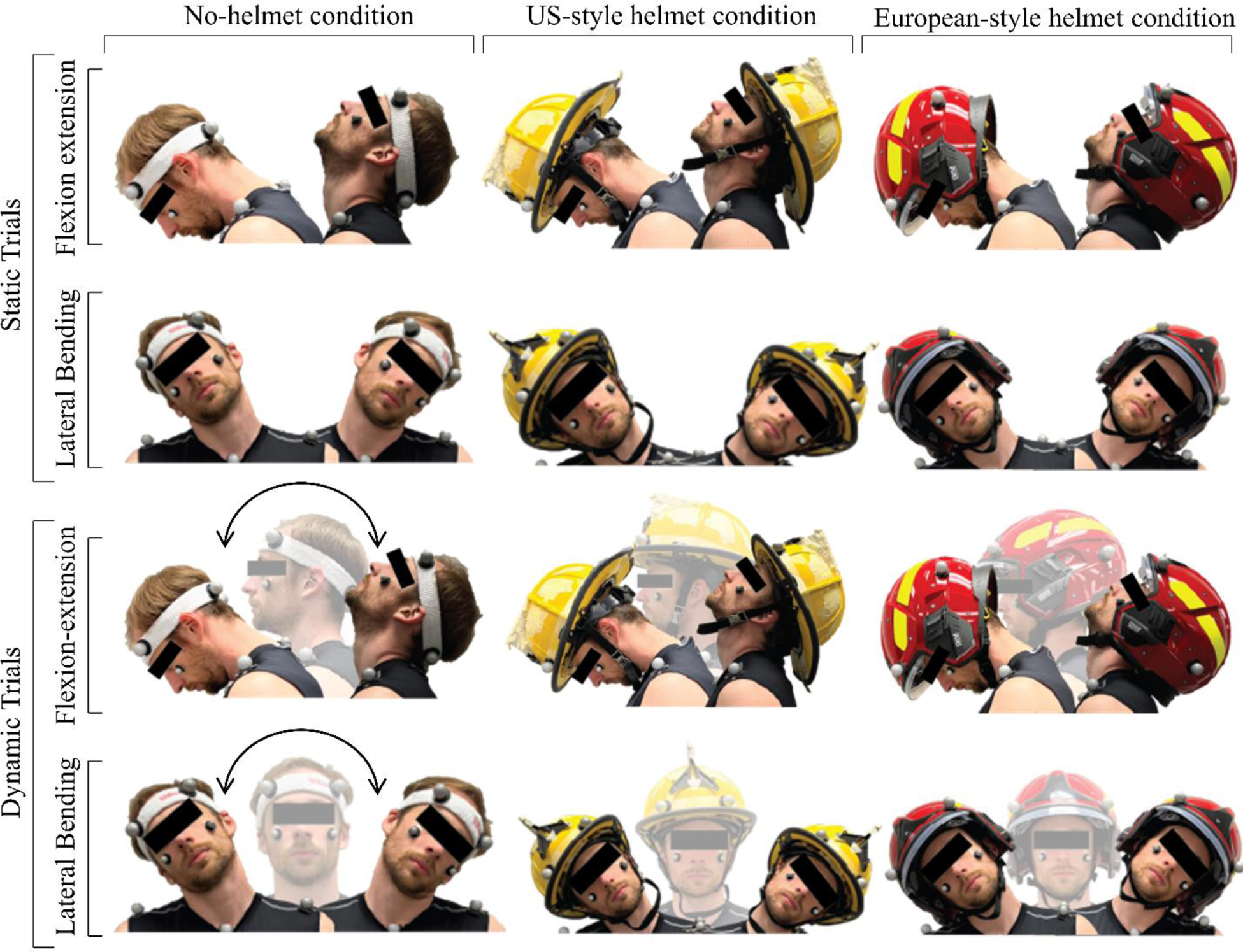
A schematic presentation of (a) Task 1 wherein subjects performing static flexion, extension, left, and right bending tasks and (b) Task 2, in which they performed dynamic flexion-extension and dynamic lateral bending tasks. Both tasks were performed for three different conditions: 1) baseline, no-helmet (left), 2) US-style helmet (middle), and 3) European-style helmet (right) in a random order.

### OpenSim Modeling

The MASI (Musculoskeletal model for the Analysis of Spinal Injury) model, a validated full-body model developed by Cazzola et al. (2017), was used as a base model for our study. Then, we added infra-hyoid muscles to the base model—taken from HYOID model (Mortensen et al., 2018)—to achieve more biofidelic cervical intervertebral kinematic results. The modified model included 35 rigid body segments, 34 body joints, 23 torque actuators, 98 Hill’s type neck muscles, and inherited head and neck models of Vasavada et al. (1998) that implements three rotational (row, pitch, and yaw) degrees of freedom (DoF) to C0-C1(atlanto-occipital), C1-C2 (atlanto-axial), C2-C3, C3-C4, C4-C5, C5-C6, C6-C7, and C7-T1 joints. We also added an additional 65 model markers in OpenSim to make the model compatible with the Cortex marker set up.

Three different versions of our modified model—a no-helmet, a US-style helmet, and a European-style helmet model—were created. Prior to the creation of head-helmet OpenSim interfacing, the geometrical shape of each helmet type was created in NMSBuilder (Valente et al., 2017) software and their inertial properties (Table 1) were estimated in ANSA (BETA CAE Systems SA, Greece) finite element pre-processor platform. It was observed that the COM of the US-style helmet is 5.8 cm superior and 1.7 cm posterior than the European-style helmet. However, the European-style helmet was 250 g heavier than the US-style helmet. Additionally, we calculated the combined moment of inertia (MOI) of head and helmet with respect to the C0-C1 joint by applying the parallel axis theorem.

**Table 1:** Inertial properties (mass, center of mass, and moment of inertial) of US-style and European-style helmets. X, Y, and Z respectively indicate anterior-posterior, inferior-superior, and medio-lateral directions. A positive value refers to anterior, right lateral, and superior directions. The center of mass was calculated with respect to the C0-C1 joint.

|  | Mass<br>(kg) | Center of Mass (m) |  |  | Moment of Inertia<br>(kg.m<sup>2</sup>) |  |  | Moment of Inertia<br>combined with the<br>head (kg.m<sup>2</sup>) |  |  |
| --- | --- | --- | --- | --- | --- | --- | --- | --- | --- | --- |
|  |  | X | Y | Z | Ixx | Iyy | Izz | Ixx | Iyy | Izz |
| US-style helmet | 1.77 | 0.002 | 0.152 | -0.006 | 0.017 | 0.028 | 0.022 | 0.092 | 0.088 | 0.10 |
| European-style helmet | 2.02 | 0.019 | 0.094 | -0.008 | 0.023 | 0.03 | 0.031 | 0.076 | 0.068 | 0.090 |

### OpenSim Simulation

Though we controlled the pace and duration of the experimental tasks using a metronome, slight discrepancies remained across the tasks and the subjects. Therefore, we down-sampled both kinematics and force plate data to 100 data points for between-task and between-subject comparisons. In OpenSim, we implemented inverse pipelines (Figure 2) that include subject-specific model scaling, inverse kinematics (IK), inverse dynamics (ID), and static optimization (SO) processes. We scaled (RMS:< 1cm; maximum: < 2 cm) each custom-made OpenSim model by using marker data collected during corresponding baseline, static head-neck neutral posture for each helmet condition. In head-helmet OpenSim models, we applied a welded joint to the head-helmet interface. We then used task-specific motion capture data to perform Inverse Kinematics (IK) simulations. In some trials, adjustments were made to the model markers and their weightages in order to keep the root mean square (RMS) < 2 cm and maximum marker errors < 4 cm during the IK analysis. The IK and the ground reaction force data were then used as inputs to perform SO. We accepted the IK results if both the SO and the OpenSim forward pipelines were successfully run and the moment and kinematic errors between forward and inverse solutions remained below acceptable thresholds.

**Figure 2:**
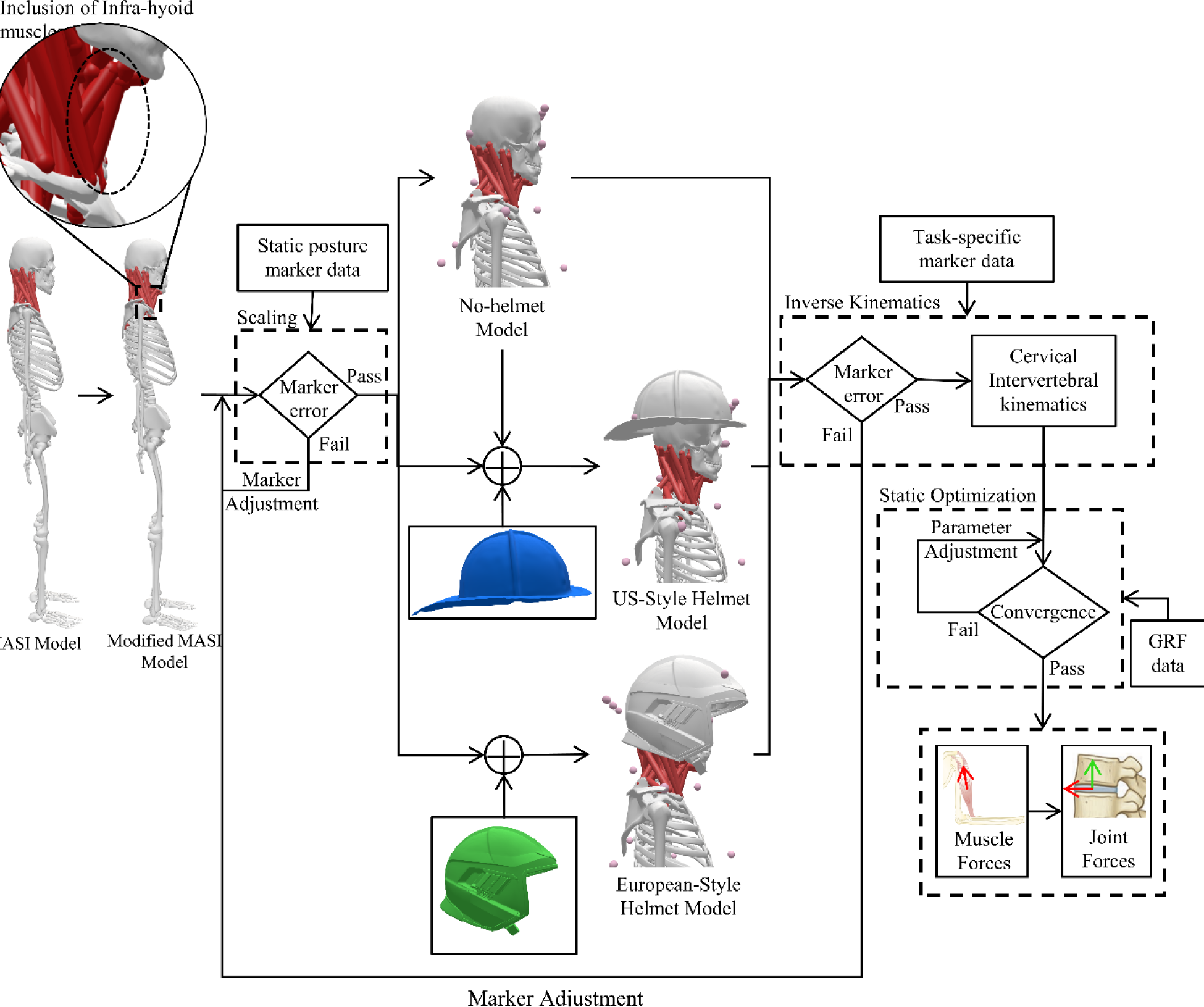
OpenSim inverse-solution workflow displaying the development of three modified MASI models: 1) no-helmet, 2) US-style helmet, and 3) European-style helmet conditions and their inverse simulation pipelines to calculate task-specific neck and cervical kinematics using experimental data.

The neck angle was calculated by summing the kinematics of all cervical intervertebral joints (C0-C1 to C7-T1). The flexion-extension and lateral bending ROM of individual cervical joints and the neck was calculated by taking the difference between their maximum positive and negative angular values during respective dynamic motions. Due primarily to poor marker (pelvis and other markers of some subjects moved or fall off during the experimental tasks) data and some challenges during experimental data collection, we abled to simulate OpenSim models of a total of 24 subjects—12 males (weight: 90.6 ± 19.1 kg; height: 1.77 ± 0.066 m; BMI: 28.8 ± 2.85 kg/m^2^; Age: 38.2 ± 7.97 years) and 12 females (weight: 67.4 ± 8.28 kg; height: 1.67 ± 0.055 m; BMI: 24.2 ± 2.85 kg/m^2^; Age: 31.2 ± 8.62 years)—and the further analysis of this study was based on these 24 best subject’s data.

### Statistical Analysis

We calculated the descriptive statistics (mean and standard error) of peak flexion, peak extension, peak left bending, peak right bending, flexion-extension ROM, and lateral bending ROM of individual cervical intervertebral joints and the neck (C0-T1) as a whole for all static and dynamic exertion trials across all subjects (males and females separately). In order to investigate the effects of helmets and sex and their interaction on neck and cervical intervertebral peak angles and ROM data, we first evaluated data normality and homoscedasticity conditions. As the majority of the peak angle data violated these two conditions even after logarithmic, exponential, and power transformations, we employed Friedman non-parametric test wherein the effects of *helmet condition* (no-helmet, US-style helmet, and European-style helmet) and *subject* were respectively treated as fixed and randomized block variables. Additionally, we considered Kruskal-Walis non-parametric test to study the *sex* (male and female) effect. In both Kruskal-Walis and Friedman tests, peak flexion, extension, left bending, and right bending angles of C0-C1, C1-C2, C2-C3, C3-C4, C4-C5, C5-C6, C6-C7, and C7-T1 joints and the neck (C0-T1) during static and dynamic tasks (task conditions) were treated as dependent variables. As the ROM data of the neck and the cervical joints met the normality assumption, we performed multi-factor Analysis of Variance (ANOVA) tests wherein *helmet condition*, *sex,* and their interaction were independent variables and the *subject* was treated as a random block. All statistical tests were performed at 95% confidence level (*α* = 0.05).

## Results

### Helmet Effect

#### Peak Neck Angle Data

Friedman tests showed that, the helmet use led to a significant increase in neck flexion (static: p<0.001, 9.55% for US-style and 5.90% for European-style; dynamic: p<0.001, 6.79% for US-style and 3.93% for European-style) and a significant decrease in neck extension (static: p=0.004, 10.7% for US-style and 2.67% for European-style; dynamic: p=0.001, 17.3% for US-style and 4.99% for European-style) in comparison to no-helmet baseline (Table 2). Particularly, the US-style helmet exhibited a larger increase in neck flexion (static: 3.45% and dynamic: 2.76%) and a greater reduction in neck extension (static: 8.23% and dynamic: 13.0%) than the European-style helmet. Though the helmet use was not significant for neck lateral bending tasks (Table 2), the use of US-style helmet led to a larger lateral bending angle (right: 2.93% and left: 4.73%) than the European-style helmet, particularly to the right side (2.50%) than the left side.

**Table 2:** Peak flexion, extension, right bending, and left bending neck angles while holding static neck postures and performing dynamic neck movements.

|  |  | Static neck posture |  | Dynamic neck movement |  |
| --- | --- | --- | --- | --- | --- |
| Peak Flexion (°) | No Helmet | Male: 33.1 ± 5.22 | 35.6 ± 1.01 | Male: 36.3 ± 4.48 | 35.62 ± 1.02 |
|  |  | Female: 38.0 ± 2.22 |  | Female: 37.7 ± 4.05 |  |
|  | US-style Helmet | Male: 38.7 ± 1.32 | 39.0 ± 0.21 | Male: 37.8 ± 3.08 | 38.04 ± 0.53 |
|  |  | Female: 39.3 ± 0.18 |  | Female: 38.2 ± 1.71 |  |
|  | European-style Helmet | Male: 37.3 ± 3.03 | 37.7 ± 0.75 | Male: 36.3 ± 4.48 | 37.02 ± 0.92 |
|  |  | Female: 38.0 ± 3.92 |  | Female: 37.7 ± 4.05 |  |
| Peak Extension (°) | No Helmet | Male: 45.6 ± 8.45 | 48.7 ± 1.75 | Male: 43.3 ± 10.8 | 45.32 ± 2.34 |
|  |  | Female: 51.7 ± 6.60 |  | Female: 47.3 ± 10.9 |  |
|  | US-style Helmet | Male: 41.0 ± 11.2 | 43.5 ± 2.02 | Male: 39.2 ± 13.6 | 37.47 ± 2.48 |
|  |  | Female: 46.0 ± 6.58 |  | Female: 35.8 ± 8.98 |  |
|  | European-style Helmet | Male: 44.8 ± 11.1 | 47.4 ± 2.00 | Male: 41.6 ± 11.4 | 43.06 ± 2.08 |
|  |  | Female: 50.0 ± 6.44 |  | Female: 44.6 ± 7.66 |  |
| Peak Right Bending (°) | No Helmet | Male: 24.6 ± 4.25 | 26.4 ± 0.94 | Male: 27.2 ± 4.02 | 28.26 ± 0.82 |
|  |  | Female: 28.2 ± 3.74 |  | Female: 29.3 ± 3.32 |  |
|  | US-style Helmet | Male: 27.3 ± 2.30 | 28.1 ± 0.58 | Male: 27.7 ± 3.30 | 27.58 ± 0.69 |
|  |  | Female: 29.0 ± 2.88 |  | Female: 27.4 ± 3.19 |  |
|  | European-style Helmet | Male: 26.8 ± 3.69 | 27.3 ± 0.73 | Male: 26.0 ± 3.33 | 26.5 ± 0.74 |
|  |  | Female: 27.8 ± 3.07 |  | Female: 27.0 ± 3.50 |  |
| Peak Left Bending (°) | No Helmet | Male: 24.8 ± 2.87 | 26.5 ± 0.77 | Male: 24.3 ± 7.49 | 26.27 ± 1.31 |
|  |  | Female: 28.2 ± 3.41 |  | Female: 28.2 ± 3.49 |  |
|  | US-style Helmet | Male: 27.4 ± 2.85 | 28.8 ± 0.63 | Male: 27.4 ± 4.21 | 28.28 ± 0.79 |
|  |  | Female: 30.2 ± 2.28 |  | Female: 29.2 ± 2.84 |  |
|  | European-style Helmet | Male: 27.2 ± 3.36 | 27.5 ± 0.73 | Male: 24.9 ± 6.10 | 26.24 ± 1.04 |
|  |  | Female: 27.9 ± 3.52 |  | Female: 27.6 ± 2.50 |  |

#### Neck ROM Data

The ANOVA results exhibited that the helmet use significantly reduced the neck flexion-extension ROM (p=0.0395), especially the US-style helmet (∼6.67%) followed by the European-style helmet (∼1%) (Table 3). While no significant effect of helmets was observed during lateral bending tasks (p=0.0693), the US-style helmet increased the lateral bending ROM by 3.13% whereas the European-style helmet decreased the lateral bending ROM by 2.76%. While compared with the experimentally measured data from the literature, our baseline, no-helmet condition data showed notably lower ROM values for both flexion-extension (14.1%∼47.1%) and lateral bending (11.6%∼54.8%) neck movements (Table 3).

**Table 3:**
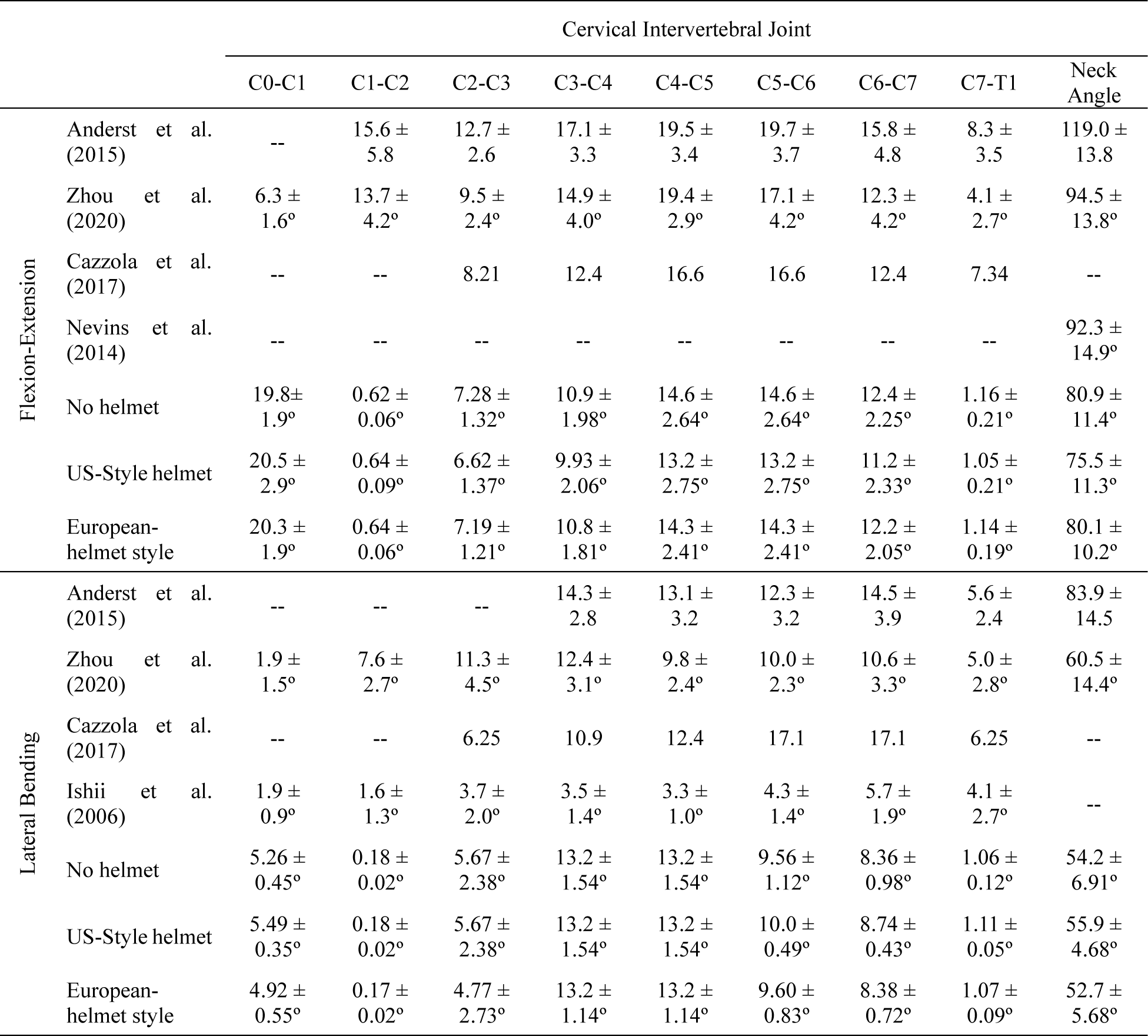
Flexion-extension and lateral bending full range of motion for each cervical intervertebral joint and the total neck (C0-T1). The total neck angle in Anderst et al. (2015) is defined as the angle between the head and the torso of the subjects.

#### Peak Cervical Intervertebral Kinematic Data

The significant effects (p<0.010) of the helmet use were observed for the majority of the intervertebral joints’ peak flexion, extension, and left and right lateral bending angles, except five scenarios – C2-C3 left and right lateral bending during static task, C2-C3 right bending during dynamic task, C0-C1 and C1-C2 extension (Figure 3). The C0-C1 joint contributed the most towards the total neck peak angles during both static (21% and 26% for neck flexion and extension, respectively) and dynamic (22% and 29% for neck flexion and extension, respectively) tasks, followed by C4-C5, C5-C6, and C6-C7 joints. Interestingly, the C0-C1 joint exhibited trivial contributions (lower than 10%) during both lateral bending motions (Figure 3).

**Figure 3:**
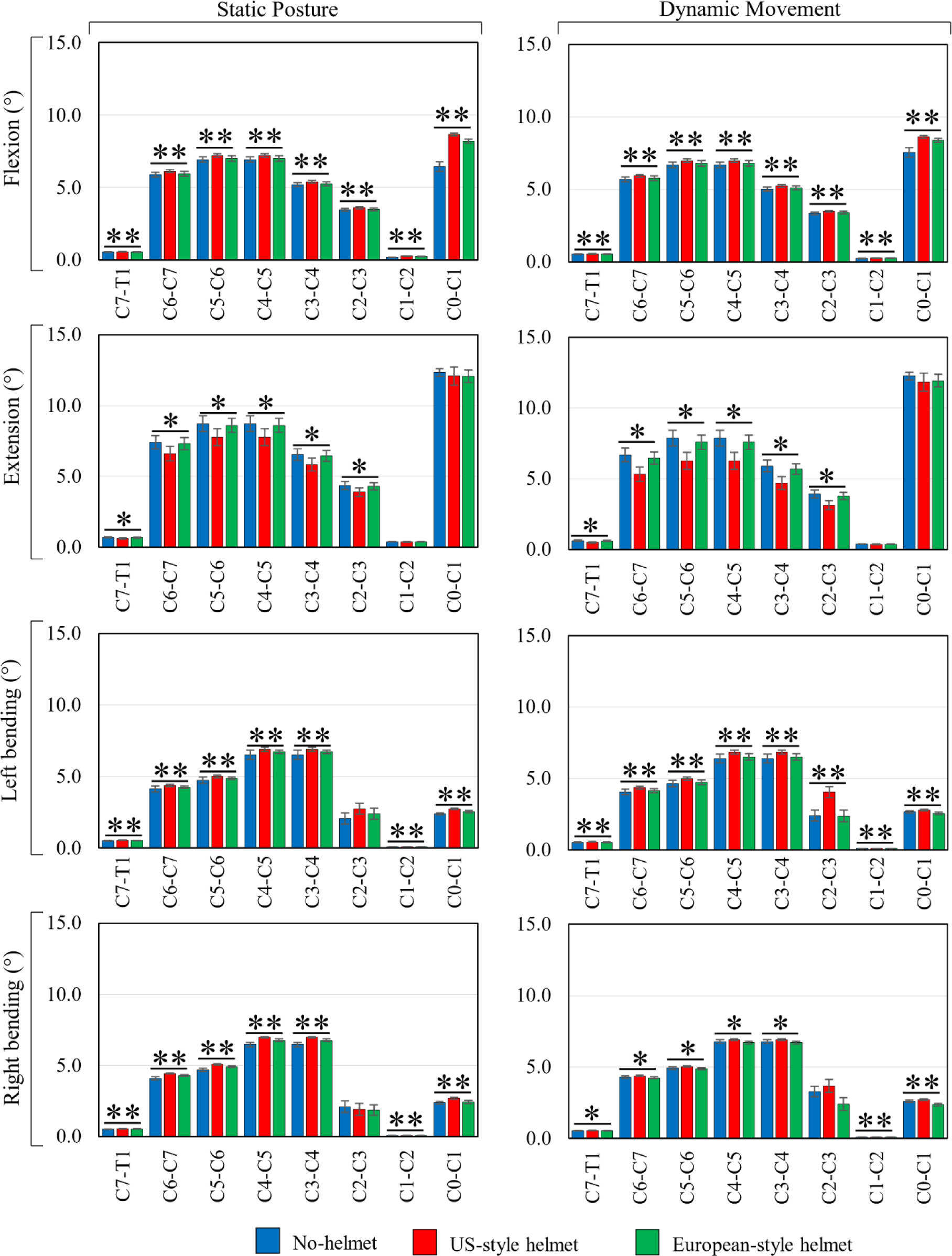
The peak angles of individual intervertebral joints (C0-C1, C1-C2, C2-C3, C3-C4, C5-C6, and C7-T1) in extreme flexion, extension, and left and right lateral bending positions for all helmet conditions: no-helmet, US-style helmet, and European-style helmet. If the helmet condition was statistically significant for a joint, it is indicated with asterisks (*: p-value < 0.05; **: p-value < 0.01) symbol.

#### Cervical Intervertebral ROM Data

The ANOVA tests displayed a significant effect of helmet use on both flexion-extension and lateral bending ROM for six cervical joints (C2-C3 to C7-T1), except C0-C1 and C1-C2 joints. Compared to the literature data, we also found noticeable discrepancies for both C0-C1 and C1-C2 joint ROM (Table 3). This discrepancy can be attributed to how the OpenSim model distributed the angle across the cervical intervertebral joints. To explore it further, we calculated the distribution ratio between the dependent joint coordinates (movement) with respect to their coupled independent joint coordinate (movement) (Table 4). We observed that motions of the lower and mid cervical joints (C2 to C7) were somewhat evenly distributed (max ratio=12.6), while the motion of the upper cervical joints was mainly carried by C0-C1 joint (max ratio=31.9). The independent joint coordinates of C1-C2 and C7-T1 remain same irrespective of intersubject variations and movement types.

**Table 4:** Ratio between dependent and independent cervical intervertebral joint. The joint mobility of C6-C7 to C2-C3 joints is dependent on the C7-T1 joint, while C0-C1 is dependent on C1-C2 joint.

| Degree of freedom<br>(DoF) | Cervical Intervertebral Joints Ratio |  |  |  |  |  |
| --- | --- | --- | --- | --- | --- | --- |
|  | C7-T1 |  |  |  |  | C1-C2 |
|  | C6-C7 | C5-C6 | C4-C5 | C3-C4 | C2-C3 | C0-C1 |
| Flexion-Extension | 10.7 | 12.6 | 12.6 | 9.45 | 6.30 | 31.9 |
| Lateral Bending | 7.85 | 9.00 | 12.4 | 12.4 | 11.2 | 28.6 |

#### Cervical Intervertebral Movement Patterns

The flexion and extension movement patterns (Figure 4) showed that subjects went into the peak flexion and extension positions, respectively, by about 2% and 5% faster (averaged across all joints) with the US-style helmet than the no-helmet condition. In contrast, the European-style helmet exhibited about 1% faster flexion and 3% slower extension movements across all joints than the no-helmet baseline. Similarly, lateral bending movements (Figure 5) of the neck and all joints (except C2-C3 joint) showed that subjects went into peak left and right bending positions, respectively, by 5% and 9% faster with the US-style helmet, followed by 4% and 7% faster with the European-style helmet, compared to the no-helmet baseline.

**Figure 4:**
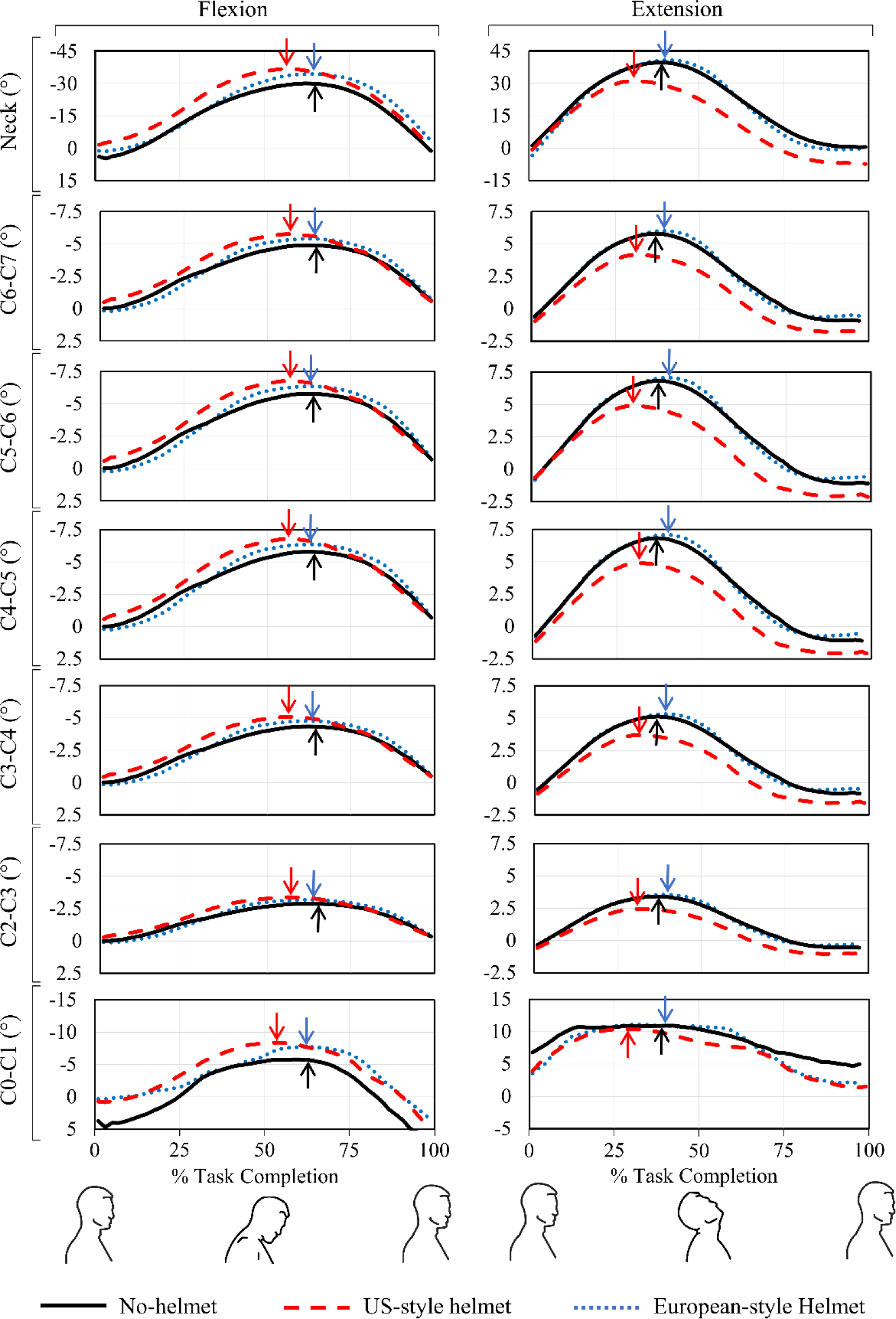
Flexion and extension movements of cervical spine joints for different helmet conditions. The C1-C2 and C7-T1 joints were not included as their movements were insignificant. The time-to-peak flexion and extension angles are indicated with arrow signs: red, blue, and black refer to corresponding time-to-peak angles for US-style helmet, European-style helmet, and no-helmet conditions, respectively. (Anderst et al., 2015a)

**Figure 5:**
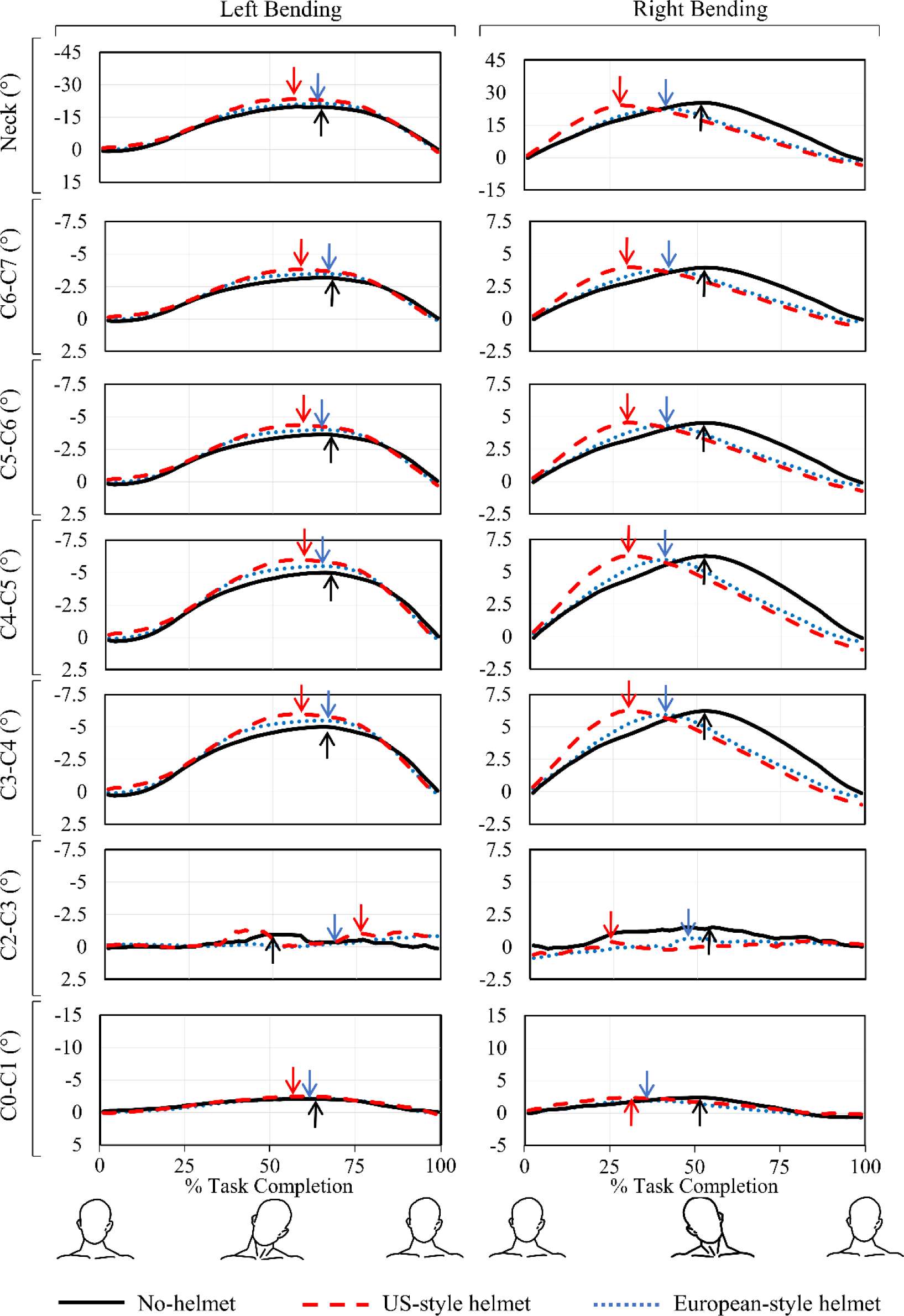
Left and right bending movements of cervical spine joints for different helmet conditions. The C1-C2 and C7-T1 joints were not included as their contributions in total neck movements were trivial. The time-to-peak flexion and extension angles are indicated with arrow signs: red, blue, and black refer to corresponding time-to-peak angle for US-style helmet, European-style helmet, and no-helmet conditions, respectively. (Anderst et al., 2015a)

### Sex Effect

The statistical tests for the sex effect did not show any statistical significance across all ROM and peak angle data of the neck and individual cervical intervertebral joints. However, females broadly exhibited greater peak flexion, peak lateral bending, flexion-extension ROM, and lateral bending ROM than their male counterparts, irrespective of the helmet use (Table 2).

## Discussion

The present study investigated the effects of firefighter helmet on cervical intervertebral kinematics during static and dynamic neck flexion, extension, right, and left bending tasks. Results demonstrated that the helmet COM had more pronounced effects on the cervical intervertebral kinematics than the helmet mass, particularly during full flexion and extension positions. In full flexion position, despite being lighter (12.4%; 250 g), US-style helmet exhibited more hyperflexion and quicker attainment of peak flexion angles than European-style helmet. Particularly, the longer moment arm of the US-style helmet in the superior direction (38.2% and 5.8 cm more superior) induced a larger rotational torques at each cervical joint and caused more accelerated dynamic neck movements than the European-style helmet. Likewise, despite having a lesser anterior COM location than the European-style helmet, the US-style helmet reached peak extension angle more rapidly. This can be attributed to its superior COM that yielded about 10% higher MOI at C0-C1 joint than those observed with European-style helmet. Similarly, though the COM of the US-style helmet was less offset in the left lateral direction than the European-style helmet, it increased the peak left bending angle by 4.51% (1.3°) because its superior COM caused about 17.4% (0.016 kg.m^2^) higher MOI in the superior direction. Thus, our findings highlighted the importance of a low-profile (i.e., less superior COM) helmet in order to yield a greater neck range of motion to the users. Finally, we observed some sex-specific insignificant differences in neck peak angles which aligns well with a previous study (Pan et al., 2018).

Interestingly, our results revealed a greater contribution from C0-C1 joint and a trivial contribution from C1-C2 joint during both flexion-extension and lateral bending movements. However, previous studies (Anderst et al., 2015b; Ishii et al., 2006; Zhou et al., 2020) showed an equal contribution (∼11°) from both C0-C1 and C1-C2 joints during neck flexion-extension movements. This discrepancy was owing to the fact that the MASI model divided the cervical spine into two independent segments: 1) upper cervical spine (C0-C2) spine with C1-C2 as an independent rotational DoF and 2) mid and lower cervical spine (C2-T1) with C7-T1 as an independent rotational DoF (Cazzola et al., 2017; Vasavada et al., 1998). The other cervical joints in each of these two segments have dependent rotational DoFs and their kinematics are estimated as a percentage of the total motion of their corresponding independent rotational DoFs. Consequently, this led to a greater contribution from C0-C1 joint and trivial movement in C1-C2 in the upper cervical spine, in addition to a very slight movement in C7-T1 joint in the mid-lower cervical spine.

Furthermore, the variations between OpenSim-derived ROM data and experimentally-measured literature data can be attributed to several factors. The experimental studies by Anderst et al. (2015b); Ishii et al. (2006); Zhou et al. (2020) considered both translational and rotational movements (six DoFs) of the cervical intervertebral joints and the intervertebral joint angles were directly measured between the adjacent vertebra’s anatomical planes. In contrast, existing OpenSim neck models employ three rotational DoFs and calculate the cervical intervertebral joint kinematics on a fixed axis of rotation (Amevo et al., 1991). Previous studies reported that the neck ROM in both flexion-extension and lateral bending decreases with age and males having significantly lower ROM than females after their 30s (Pan et al., 2018). Thus, our findings of reduced neck and cervical intervertebral mobility can be associated to the fact that our study subjects were comparatively older and bulkier (high BMI) than reported experimental studies.

This study has several limitations. First, the effects of helmet inertial properties on neck rotation were not evaluated. Second, we recruited and analyzed a limited number of subjects and two different versions of firefighter helmets. Third, a certain extent of variations in estimating the neck and cervical intervertebral angles can be associated to the OpenSim optimization routines of scaling and IK processes; these variations would be different if we used a different modeling platform. Fourth, as existing OpenSim neck models do not include translational DoFs and measure intervertebral rotational kinematics by employing a fixed proportion, our data on cervical kinematics are an estimated representation of the actual cervical intervertebral movement. In summary, the present study is a first-of-its-kind investigation on the influence of helmet inertial properties on the cervical intervertebral kinematics during both static and dynamic neck exertions. Our findings established the critical role played by the helmet COM in adversely affecting the cervical spinal mobility and acknowledged the placement of helmet COM near to C0-C1 joint (i.e., designing a low-profile helmet) in order to reduce potential neck injuries during prolonged wear.

## Acknowledgements

This work was partly supported by the U.S. Department of Homeland Security (70RSAT21CB0000023). We would like to acknowledge Mr. Felipe S. Zambrini, Mr. Leonardo Wei, Mr. Hossein Bahreinizad, and Ms. Kathryn Bell for their assistance in data collection and data preprocessing.

